# Characterizing and controlling nanoscale self-assembly of suckerin-12

**DOI:** 10.1101/2020.08.10.244673

**Authors:** Jasmine M. Hershewe, William D. Wiseman, James E. Kath, Chelsea C. Buck, Maneesh K. Gupta, Patrick B. Dennis, Rajesh R. Naik, Michael C. Jewett

## Abstract

Structural proteins such as the “suckerins” present promising avenues for fabricating functional materials. Suckerins are a family of naturally occurring block copolymer-type proteins that comprise the sucker ring teeth of cephalopods and are known to self-assemble into supramolecular networks of nanoconfined *β*-sheets. Here, we report characterization and controllable, nanoscale self-assembly of suckerin-12 (S12). We characterize impacts of salt, pH, and protein concentration on S12 solubility, secondary structure, and self-assembly. In doing so, we identify conditions for fabricating ~100 nm nanoassemblies (NAs) with narrow size distributions. Finally, by installing a non-canonical amino acid (ncAA) into S12, we demonstrate the assembly of NAs that are covalently conjugated with a hydrophobic fluorophore, and the ability to change self-assembly and *β*-sheet content by PEGylation. This work presents new insights into the biochemistry of suckerin-12 and demonstrates how ncAAs can be used to expedite and fine-tune the design of protein materials.

## Introduction

Self-assembling and stimulus-responsive proteins hold promise for engineering functional materials and nanomedicines.^1–6^ With exact sequence control, stimuli responsiveness, and biocompatibility, block-copolymer proteins, such as elastin-like polypeptides, silk fibroins, and the recently-discovered “suckerins” are of particular interest for fabricating materials.^7–15^ Additionally, the ability to incorporate non-canonical amino acids (ncAAs) into protein materials has further expanded their functionality.^16–22^ The suckerins are a family of structural proteins that make up the sucker ring teeth (SRT), a circular structure with hook-like teeth located in the arms of cephalopods that assist in wrangling prey.^13^ With an elastic modulus on the order of GPa, SRT rival synthetic polymers in strength despite the absence of nature’s common hardening methods like biomineralization and crosslinking.^23^ Rather, the mechanical strength of SRT is derived from weak protein-protein interactions that stabilize a semi-crystalline protein mesh that is mechanically reinforced by precisely-sized, nanoscale *β*-sheets.^13,24,25^ Suckerins have conserved, modular, di-block copolymer architectures that consist of *β*-sheet domains flanked by aromatic-rich amorphous domains. This architecture is intimately linked to the emergent supramolecular assembly of suckerins, permitting (i) formation of H-bond networks with inter- and intra-molecular *β*-sheets, and (ii) π-π stacking of aromatic residues in distally-located amorphous domains.^26,27^

While the modular architecture of suckerins plays key roles in assembly, strength, and processing of natural SRT, the structural and physiochemical properties of recombinant suckerin proteins and peptides have been exploited to make synthetic materials.^11,27,28^ Building upon a robust field of work with *β*-sheet rich silk fibroins, Miserez and others have pioneered the design of suckerin-based functional materials, including substrates for metallic nanoparticle growth,^29^ underwater adhesives,^18^ and crosslinked hydrogels.^30,31^ Of key importance for the present work, the Histidine (His)-rich 39-kDa suckerin-19 (S19) was recently shown to assemble into stable, *β*-sheet rich nanoassemblies (NAs).^32^ This work established that S19, which naturally forms polydisperse colloids in aqueous solution upon purification, can be controllably assembled into relatively monodispersed NAs of a range of sizes.^32,33^ Self-assembly was achieved using a salting-out method where simple manipulation of solvent pH and ionic strength led to controllable chain collapse, *β*-sheet formation, and subsequent self-assembly of NAs. The densely packed *β*-sheets in S19 NAs enabled noncovalent encapsulation of DNA and hydrophobic small molecules, while the high His content enabled pH-dependent intracellular cargo release in mouse models.^32^ These results set precedence for how the unique features of suckerins can be leveraged to fabricate functional nanocarriers for biomedical applications or in other functional materials.

As crucial parameters in the engineering of nanocarriers,^34^ we sought to access new suckerin NA particle sizes and compositions in this work. Suckerin proteins range in size (~5-60 kDa), isoelectric point (pI ~7-10), relative proportion and spacing of repeat domains, and the number of repeat units within the protein sequence.^24^ Since the amino acid sequence dictates the stability, geometry, and presentation of *β*-sheets, which are critical for self-assembly, the family of suckerins are a potential toolbox for making NAs with tunable sizes and pH-responses.

In this study, we report biochemical characterization and controllable self-assembly of the 23-kDa suckerin-12 (S12). S12 has a similar architecture and sequence to S19, including a His-rich primary sequence, yet is a smaller, more soluble protein. With an eye toward controllable self-assembly, we manipulate solvent conditions to controllably unfold and refold *β*-sheets in S12. By varying salting-out process conditions (pH, salt type, protein concentration, and ionic strength), we identify conditions for salting-out ~100 nm NAs with narrow size distributions. Additionally, we show that S12 can be efficiently expressed and purified with a non-canonical amino acid (ncAA) handle for site-specific, biorthogonal functionalization. We use the ncAA to covalently pre-functionalize S12 with a small, hydrophobic fluorophore before self-assembly, resulting in fluorophore-loaded NAs without the need to change self-assembly conditions. Finally, we show that altering the hydrophobicity of S12 through PEGylation changes self-assembly and *β*-sheet enrichment. Taken together, this work presents new biochemical characterizations of S12, and shows the utility of ncAAs to accelerate building and testing new protein materials.

## Experimental (Materials and Methods)

### Expression of S12wt and S12-pAzF

The open reading frame of S12 was obtained from Uniprot (Identifier # A0A075LXT7_DOSGI) and synthesized by Integrated DNA technologies without the signal peptide. The gblock was cloned into the IPTG-inducible pET28a (Novagen) to make the expression plasmid pET28a-S12. For S12-pAzF, the codon encoding Glycine 32 was mutated to the amber stop codon (TAG) via PCR mutagenesis, resulting in the pET28a-S12-TAG expression vector. For S12wt, pET28a-S12 was transformed into BL21 Star (DE3), selected on LB agar plates supplemented with 50 ug/mL Kanamycin, and grown overnight at 37 °C. For S12-pAzF, pET28a-S12-TAG was co-transformed with the arabinose-inducible pEVOL-pAzFRS.2.t1 (Addgene, Plasmid #73546) and selection plates contained 50 μg/mL Kanamycin and 34 μg/mL Chloramphenicol. Cells were always freshly transformed. For expression, single colonies were selected and grown overnight at 37 °C in 2xYTP media supplemented with appropriate antibiotic(s). Expression cultures were grown in shake flasks (1 L scale) or a Sartorius Stedim BIOSTAT Cplus bioreactor (10 L scale) at 37 °C with antibiotics. S12 expression was induced during exponential growth at OD 0.6-0.8 with 0.5 mM IPTG. For S12-pAzF expression, cultures were supplemented with 0.5 mM IPTG, 0.1% (w/v) arabinose, and 3 mM pAzF (Chem-Implex Int’l Inc.) at induction. Expression proceeded overnight at 30 °C. Expression cultures were harvested at 5,000 *x g* for 15 mins at room temperature. Cells were washed once with Buffer 1 (50 mM Tris, pH 8 and 100 mM NaCl), flash frozen with liquid nitrogen, and stored at −20 °C until purification.

### Inclusion body purification of S12 and S12-pAzF

For purification, we adapted an inclusion body preparation from Buck and colleagues.^30^ Unless otherwise noted, all steps were performed at room temperature. Cell pellets were thawed and resuspended in 30 mL of Buffer 1 per 0.5 L of expression culture. Lysozyme was added to the resuspended cells to a final concentration of 1 mg/mL and incubated for 30 mins. Cells were sonicated on ice using a Q125 Sonicator (Qsonica, Newtown) with a 3.175 mm diameter probe at a frequency of 20 kHz and 45% amplitude. Energy was delivered to cells in pulses of 15 s followed by 15 s off, for 10 mins total. Lysed cells were centrifuged at 10,000 *x g* for 10 mins. Supernatant containing soluble cellular proteins was discarded and inclusion bodies containing S12 were collected in the pellet. Pellets were resuspended for washing in 35 mL of Buffer 1 supplemented with 1% Triton X-100 (v/v) by light vortexing. After dispersion, suspensions were incubated on an orbital shaker for 10 mins at 4 °C. Pellets were collected by centrifugation at 5,000 *x g* for 15 mins. Washes with Triton X-100 supplemented buffer were repeated for three total washes. After washing, pellets were resuspended in 15 mL of deionized water per 1 L of expression culture, and pellets were dispersed by light vortexing and sonication. For ‘pH-cycling’ the S12 suspension was poured into a clean beaker with stirring and pH monitoring. HCl was used to adjust the solution pH to 3 to solubilize S12. Soluble S12 was incubated for 1 hr at room temperature with constant agitation, then clarified at 20,000 *x g* for 10 mins. To precipitate S12, 1 M NaCl was mixed with the supernatant to 100 mM. Under agitation, the pH was adjusted to 8 with 1 M Tris pH 8. Insoluble S12 was recovered by centrifugation at 10,000 *x g* for 10 mins. Another round of solubilization and precipitation was conducted to improve purity. Pellets were resuspended to ~5 mg/mL of S12 in 5% (v/v) acetic acid and dialyzed exhaustively against water using 20,000 MWCO membranes (ThermoFisher Scientific, USA). For quantitation of protein concentration, the sample absorbance at 280 nm was measured using a Nanodrop 2000 (Thermo Scientific, USA). The molar extinction coefficient and molecular weight of S12 were used to calculate protein concentration. S12wt and S12-pAzF were aliquoted, flash frozen on liquid nitrogen, and stored at ~6 mg/mL at −80 °C until use.

### Strain promoted azide-alkyne cycloaddition (SPAAC) reactions

S12-pAzF was labeled using the strain promoted azide-alkyne cycloaddition (SPAAC) using various dibenzocyclooctyne (DBCO) probes. SPAAC reactions were conducted in water (unless otherwise noted) at 0.02 mM S12-pAzF incubated with 2 mM of strained alkyne probes. Reactions were incubated overnight at 30° C in an end-over-end mixer with constant rotation, followed by exhaustive dialysis into water using 20,000 MWCO membranes (ThermoFisher Scientific). To make stock solutions, DBCO-TAMRA (Millipore Sigma, USA) was resuspended to 10 mM in 100% DMSO, and DBCO-mPEG 5 kDa/ DBCO-mPEG 10 kDa (Broad Pharm) were resuspended to 10 mM in water. All DBCO probe stocks were stored at −20 °C. Labeling efficiency was determined using SDS-PAGE, followed by densitometry analysis and in-gel fluorescence.

### Turbidity profiles

The salt-response of S12-pAzF was investigated using turbidity profiles. All salts were purchased from Sigma. Solutions of the following salts were prepared at 500 mM: potassium acetate, sodium acetate, potassium chloride, sodium chloride, potassium sulfate, sodium sulfate, potassium citrate, sodium citrate. After dissolving, salts were brought to the desired pH using glacial acetic acid (regardless of the salt anion), NaOH for sodium salts, and KOH for potassium salts. 20 μL of S12-pAzF at 0.5 mg/mL in water was pipetted into the wells of a black Costar 384 well plate with clear bottom for imaging (Corning). 20 μL of salt solutions were added to each well to 250 mM salt (40 μL total volume) and mixed well. Plates were quickly spun down to remove bubbles, then sealed and equilibrated for 1 hr at room temperature on an orbital shaker. After incubation, optical density at 350 nm (OD 350) and pH were measured using a Synergy H1 microplate reader (Biotek) and an Orion pH probe (Millipore Sigma), respectively. OD 350 of triplicate wells for each condition were averaged and background subtracted by the average OD 350 of triplicate wells containing 40 μL of pH-adjusted salt at 250 mM with no protein.

### Salting-out S12 NAs

A fresh aliquot of S12-pAzF was thawed on ice and diluted to 2x of the final desired concentration in ice-cold 5% (v/v) acetic acid. To disperse naturally occurring S12 colloids, protein was sonicated for 20 mins in a cold water bath, then sterile filtered using a 0.1 μm filter (Millex-VV Syringe Filter, Merck Millipore Ltd). The resulting mixture was spun at 15,000 *x g* for 10 mins at 4 °C; the supernatant of this spin was the working stock of protein and was prepared fresh for each experiment. For salting-out NAs, cold KCl solution at 500 mM was quickly mixed 1:1 with the working protein stock and vigorously pipette mixed on ice. Final concentrations of KCl and acetic acid were 250 mM and 2.5% (v/v), respectively, unless otherwise noted. Notably, for initial optimization experiments (**Figs. S4-S5**), all steps were the same as described above, except that particles and reagents were equilibrated to room temperature before salting-out, and the concentration of protein, KCl, and acetic acid were varied as indicated.

### Fluorescence microscopy

Fluorescent NAs fabricated using the TAMRA-conjugated S12 scaffold (S12-TAMRA) were observed using fluorescence microscopy. To make TAMRA-loaded S12 NAs, S12-TAMRA at 0.22 mg/mL in 5% (v/v) acetic acid was mixed 1:1 with 500 mM KCl and pipette mixed vigorously on ice. 2 μL of sample was mounted on a glass slide for fluorescence microscopy. Particles were viewed using a Nikon Ni-U upright microscope with a 100×1.45 n.a. plan apochromat objective. Images were captured using an Andor Clara-Lite digital camera (Andor Technology) and Nikon NIS Elements software (Nikon Instruments Inc.). Fluorescence images were collected using a yellow excitation Y-2E/C filter combination.

### SDS PAGE and in-gel fluorescence assays

SDS-PAGE was run using NuPAGE 4-12% Bis-Tris protein gels with MOPS-SDS buffer (Thermo Fisher Scientific) and either the Chameleon 800 or Chameleon Duo protein standards (LI-COR Biosciences). For total protein imaging, gels were stained using SimplyBlue SafeStain Coomassie (Thermo Fisher Scientific) and imaged using a Gel Doc XR (Bio-Rad). Densitometry analysis for purity was conducted using Image J. For in-gel fluorescence assays, gels were imaged using a LI-COR Odyssey Fc (LI-COR Biosciences).

### Determination of protein solubility

To assess changes in the solubility parameters of S12-PEG conjugates against unconjugated S12-pAzF, we exposed 1 mg/mL PEGylated or unconjugated S12-pAzF to 50 mM Tris pH 8 and 50 mM MES pH 6.6. Triplicate samples were incubated for 5 mins, then clarified at 20,000 *x g* for 10 mins. The absorbance at 280 nm (A280) of the supernatant was then measured, and the soluble fraction calculated.

### Liquid chromatography mass spectrometry

Purified proteins (10 pmol, or 10 μL of 1 μM sample, each) were separated with an XBridge BEH C4 analytical column (300Å, 3.5 mm, 2.1 mm × 50 mm, SKU 186004498; Waters) fitted with a matched guard column (2.1 mm × 10 mm, SKU 186007230; Waters, USA) using an Agilent 1200 HPLC system and analyzed with an Agilent 6210A time-of-flight mass spectrometer equipped with an ESI source (Agilent). The aqueous phase (A) was 95% water 5% acetonitrile with 0.1% formic acid and the organic phase (B) was 100% acetonitrile with 0.1% formic acid and the method run at a constant 0.4 mL/min was, in brief: diversion to waste at 15.8% B for one minute; elution into source with a 12 minute gradient from 15.8% to 65.8% B; column wash with a two minute gradient from 65.8% to 69.9% B, then two mins at 100% B; column equilibration for six mins at 15.8% B. TOF-MS spectra were processed using the Agilent Mass Hunter (vB.04.00). The maximum entropy method was used to deconvolute spectra for m/z = 700 through 2,000. Deconvoluted spectra were manually analyzed for the presence/absence and relative abundances of the following species based on theoretical molecular weights: full length protein with pAzF (TAG construct) or without (wild-type construct); truncations at TAG codon; pAzF substitution with tyrosine or glycine; any protein sequence ± initiator methionine; and additional contaminants.

### Circular dichroism (CD)

CD spectra were collected in triplicate scans at 25 °C using a Jasco J-810 spectrometer using the following parameters: wavelength range of 190−260 nm, scan rate of 50 nm/min, and a bandwidth of 1 nm. Background absorbance from each buffer/pathlength combination was measured before acquiring spectra and automatically subtracted by the CDPro software during acquisition (Jasco). The spectra were averaged and smoothed using the Savitzky-Golay method using CDPro. Traces were converted to mean residue ellipticity (MRE) where appropriate. For CD measurements where pH and salt content were varied, 0.5% (v/v) acetic acid and/or 50 mM KCl was used. For CD measurements comparing PEGylated S12 (S12-5K) and unconjugated S12-pAzF, measurements were conducted with 0.1 mg/mL S12-pAzF or S12-5K in water.

### Dynamic Light Scattering (DLS) measurements

DLS measurements were performed on a Malvern Zetasizer Nano ZS with a measurement angle of 173° in disposable cuvettes with a sample volume of 70 μL (Malvern Panalytical). All measurements were collected in triplicate for 13 scans per measurement. Refractive index and viscosity were obtained from the instrument’s parameter library. The instrument’s ‘General Purpose’ setting was used to calculate particle size distributions (PSDs). Samples were analyzed directly without dilution.

Nanoparticle Tracking Analysis measurements were performed on a Malvern Nanosight NS300 using a 642 nm red laser (Malvern Panalytical). Samples were diluted directly before analysis to manufacturer-recommended particle concentrations in sterile 250 mM KCl, 2.5% acetic acid. Samples were flowed into the cell, and the instrument was focused according to manufacturer recommendations. Measurements were collected at room temperature, using a 1 mL syringe and a syringe pump infusion rate of 40 (arbitrary units). Data for each sample was collected in 3 separate 1 min videos, under continuous flow conditions. Mean particle diameters and PSDs were obtained from aggregate Nanosight experiment reports of each run, then averaged across triplicates and corrected for dilution factor. For PSDs, curves were normalized before plotting.

### Transmission electron microscopy (TEM)

NAs were visualized by TEM. For TEM measurement, 200 mesh Cu grids with a Formvar/ Silicon Monoxide (Cat #1830, Ted Pella Inc.) were placed in a Pelco easiGlow glow discharger (Ted Pella Inc.) and an atmosphere plasma was introduced on the surface of the grids for 30 seconds with a current of 15 mA at a pressure of 0.24 mbar. This treatment creates a negative charge on the carbon membrane, allowing for liquid samples to spread evenly over the grid. 4 μL of sample were pipetted onto the grid and incubated for 2 mins before wicking excess liquid from the grids. Grids were immediately washed with water, dried, and imaged. Notably, stain was not used due to sample aggregation upon staining. Samples were viewed using a Hitachi HT7700 (Hitachi, Ltd.) at 80 keV. Image data was collected by a Gatan Orius SC1000A CCD camera (Gatan Inc.). Image analysis was done with Image J.

## Results and Discussion

### Expression, characterization, and biorthogonal functionalization of S12

The block copolymer architecture and amino acid sequence of S12 are shown in **Fig. 1A**, highlighting several important biochemical features. S12 is comprised of 6 repeats of *β*-sheet forming blocks flanked by amorphous blocks. *β*-sheet blocks are Alanine-rich, resembling *β*-sheet-forming domains in silk fibroin,^35^ and amorphous blocks are Glycine (Gly) and Tyrosine (Tyr) rich. Importantly for controlled assembly and mechanical strength, *β*-sheet regions are flanked by Proline residues, confining *β*-sheets to specific nanoscale dimensions.^27^ The high Tyr content of amorphous blocks (15 % by mol) plays an important role in the supramolecular arrangement of suckerins via aromatic π-π stacking interactions that orient and stabilize distal amorphous domains. From a practical perspective, the high Tyr content also enables inter-protein di-Tyr crosslinking of *β*-sheet enriched monomers for stabilizing nanocarriers for downstream use.^30,32^ S12 is hydrophobic in character but contains 9 mol% His. Protonation of His residues is thus important for solubility, conferring high solubility of up to 60 mg/mL in slightly acidic conditions (~pH 5).^30^

**Figure 1.**
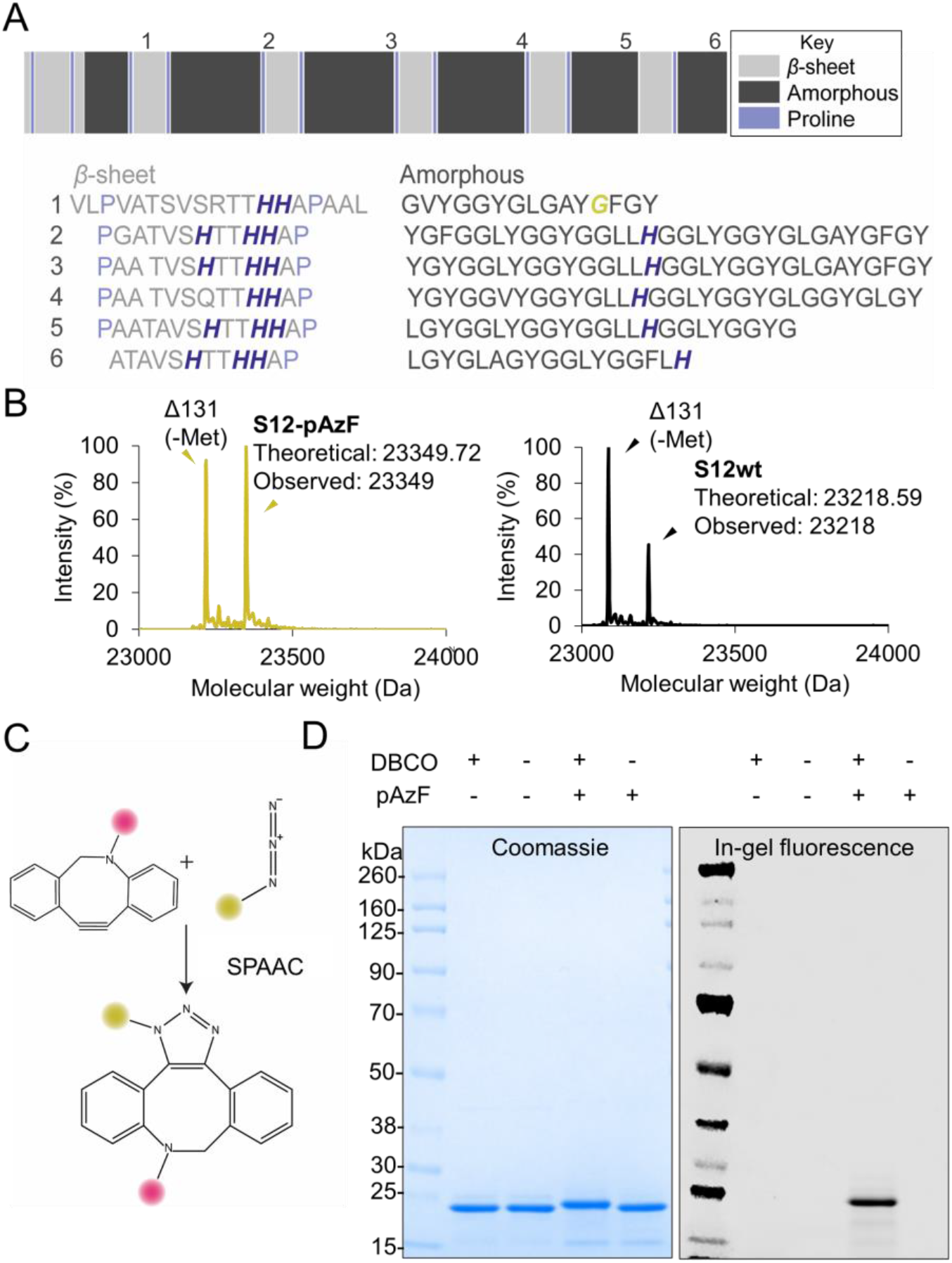
Expression, characterization, and biorthogonal functionalization of S12. **(A)** Schematic of block copolymer architecture of S12 **(*top*)**. Left to right is N- to C-terminus. Amino acid sequence of S12, with the six repeats of *β*-sheet and amorphous blocks shown **(*bottom*)**. Histidine residues are colored in dark purple. Gly 32, which was mutated to pAzF (S12-pAzF), is highlighted in gold. **(B)** Deconvoluted liquid chromatography mass spectrometry spectra of S1-pAzF **(*left*)** and S12wt **(*right*)**. Mass signatures corresponding with methionine-cleaved proteins are indicated. **(C)** The strain promoted azide alkyne cycloaddition (SPAAC) for biorthogonal coupling. The azido protein is represented as a gold circle and the DBCO-labeled partner represented as a pink circle. **(D)** SDS-PAGE analysis of protein purity and SPAAC reactions. Coomassie stain is shown ***left*** and in-gel fluorescence of the DBCO-TAMRA probe conjugated to S12 is shown ***right***. −pAzF and +pAzF refer to S12wt and S12-pAzF, respectively. Data are representative of three independent experiments.

We overexpressed and purified wild-type S12 (S12wt), and a point mutant of S12 bearing a single non-canonical amino acid (ncAA) from *Escherichia coli*-based cultures. To enable biorthogonal functionalization, we replaced the native Gly residue at position 32 with the ncAA para*-L*-azido-phenylalanine (pAzF), referred to as S12-pAzF. S12-pAzF was specifically designed such that interactions impacting pH-responsiveness and secondary structure would not be interrupted and so that subsequent biochemical studies would mimic the behavior of S12wt as closely as possible. Unless otherwise noted, all studies were performed with S12-pAzF. For purification of S12wt and S12-pAzF, we optimized a columnless pH-cycling method^30,33^ to obtain product at >90% purity (**Fig. S1**). We confirmed the molecular weights of S12 and S12-pAzF using LC-MS. The deconvoluted spectra showed two major peaks in each sample: one corresponding to the theoretical masses of the full-length product, and the other corresponding with the methionine-cleaved proteins (**Fig. 1B**). We did not observe ions corresponding with truncated or mis-incorporation products in the S12-pAzF sample, demonstrating that small truncation products do not co-purify with S12-pAzF and that pAzF-incorporation is high-fidelity.^36^ With S12-pAzF in hand, we tested for biorthogonal conjugation using the strain-promoted azide-alkyne cycloaddition reaction (SPAAC) with a strained alkyne dibenzocyclooctyne TAMRA fluorophore (DBCO-TAMRA) (**Fig. 1C**).^37^ Using SDS-PAGE to analyze SPAAC reaction products, we observed a uniform increase in molecular weight that was accompanied by an in-gel fluorescence signal, confirming the covalent conjugation of TAMRA to S12 (S12-TAMRA) (**Fig 1D**). DBCO-TAMRA did not react with S12wt, indicating that SPAAC labeling was biorthogonal and specific.

These results demonstrate several useful features of S12 and S12-pAzF. First is the ease of upstream expression and downstream purification of S12 and mutants thereof. We observed strong overexpression bands for S12, indicating that it can be produced at high yields in *E. coli* without the need for extensive optimizations (**Fig. S1**). For downstream processing of S12wt and S12-pAzF, our columnless purification protocol readily scaled to process up to 10L of expression culture and yielded >300 mg of purified protein per L culture. Yields of S12 are an order of magnitude above what has been reported for other recombinantly-produced suckerins, showing promise for scalability.^33^ We obtained purities of 98% and 92% for S12wt and S12-pAzF (**Fig. 1D**),respectively, demonstrating that, while columnless purification can be more cost-effective than traditional column chromatography, the product purity is comparable. Our results also show that orthogonal translation systems (OTSs) provide an avenue for obtaining workable yields of pAzF-incorporated proteins. Of specific importance for suckerin materials and other cationic proteins, SPAAC readily proceeded under acidic conditions, providing an avenue for functionalizing proteins that are not soluble at neutral pH conditions (**Fig. S2**). Thus, pAzF represents an option for expanding the chemistry of proteins when other common protein conjugation methods, like maleimide labeling on cysteine or N-hydroxysuccinimide ester labeling with lysine, are inaccessible due to pH constraints. It is notable that many *β*-sheet rich silk fibroins are not amenable to recombinant expression, therefore interfacing ncAA with suckerins is an attractive avenue for producing *β*-sheet rich materials that can be readily, site-specifically functionalized.^30^

### pH, salt, and protein concentration impact secondary structure and solubility

Understanding how simple biochemical parameters like protein concentration, pH, and salt impact secondary structure is foundational to engineering NAs with new suckerin isoforms. To discover the impact of these parameters on secondary structure and *β*-sheet formation in S12, we used circular dichroism (CD) spectroscopy. Since standard CD analysis software did not yield reasonable secondary structure predictions—an observation that has been made with CD analysis of other suckerin proteins—we analyzed the data qualitatively using spectral signatures that have been confirmed with Fourier-transform infrared (FTIR) spectroscopy elsewhere.^18,33^ To assess the impacts of molecular crowding on secondary structure, we obtained CD spectra at three concentrations: 0.2 mg/mL, 1.2 mg/mL, and 2.1 mg/mL (**Fig. 2A**). Protein at 0.2 mg/mL exhibited both random coil structure and *β*-sheet structure, indicated by minima at 195-205 nm and 210-220 nm, respectively. As the concentration of protein was increased, the compositions of *β*-sheet and random coil changed, with *β*-sheet eventually becoming the dominant spectral feature at the highest concentration tested.

**Figure 2.**
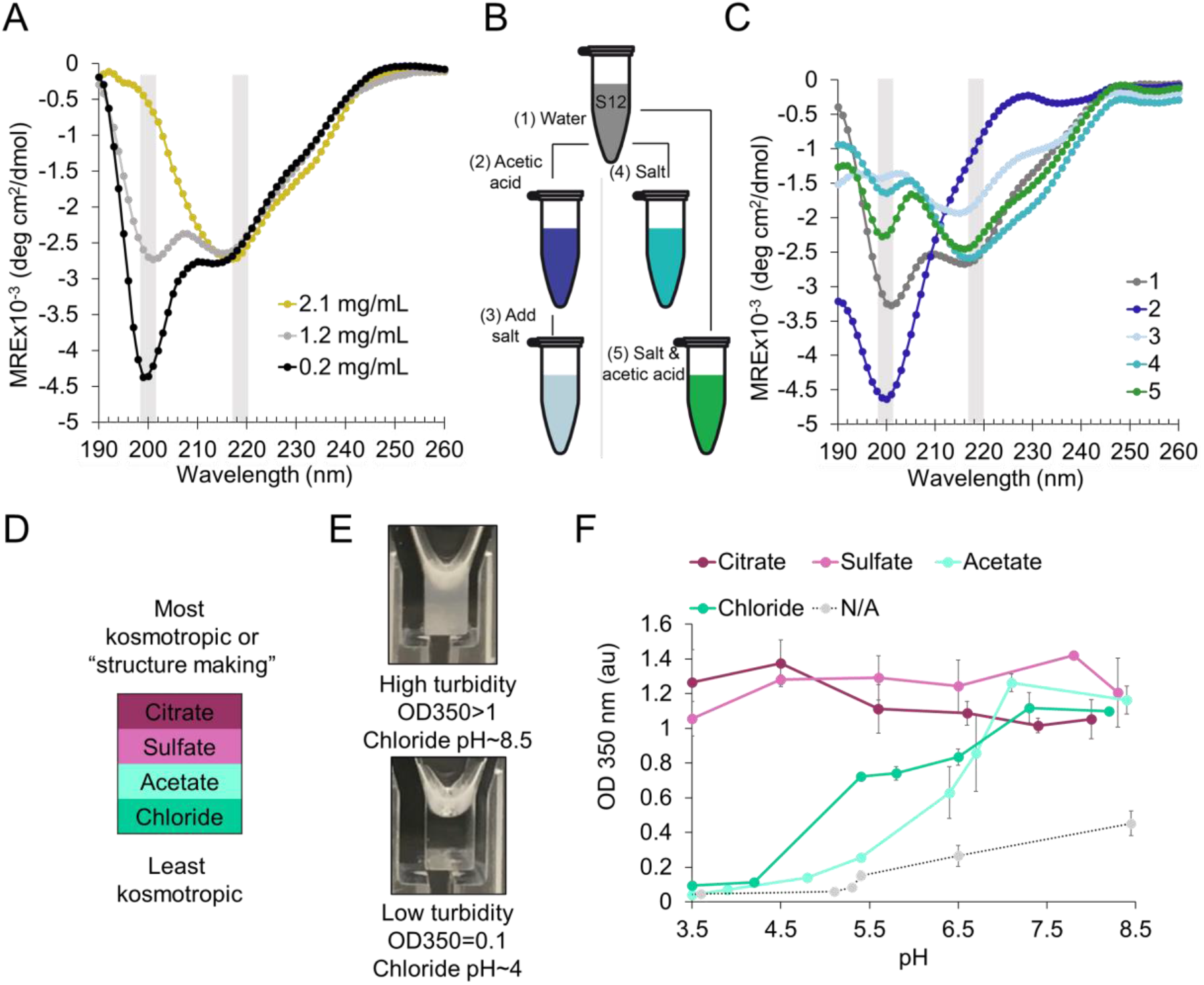
pH, salt, and protein concentration impact secondary structure and solubility. **(A)** CD spectra of S12-pAzF at a range of protein concentrations. Gray rectangles highlight the wavelength ranges corresponding with random coil (~200 nm) and beta sheet (~217-220 nm) signatures. Mean residue ellipticity (MRE) is presented. Traces represent the average of triplicate background subtracted scans. **(B)** Schematic of which components were added to each sample, and the order the components were added in. Samples are color coded to correspond with **(C)**. **(C)** CD spectra of S12-pAzF exposed to combinations of acetic acid and/or KCl. Traces represent the average of triplicate background subtracted scans. **(D)** Hofmeister series ordering of the anions tested. Anions are ordered from most kosmotropic in pink shades and least kosmotropic in green shades. **(E)** Photographs of high ***(top)*** and low ***(bottom)*** OD 350 nm samples of S12, showing how absorbance corresponds with turbidity in chloride salt. The concentration of protein both samples is equivalent. **(F)** Turbidity profiles of S12-pAzF as functions of pH and anion type. Error bars represent the standard deviation between three independent replicates.

To test impacts of pH and salt, we exposed 0.1 mg/mL solutions of S12-pAzF to acetic acid and potassium chloride (KCl), adding these components alone, together, or sequentially (**Fig. 2B**). Acidification of S12-pAzF in water resulted in a conformational switch, causing the protein to adopt predominantly random coil structure (**Fig. 2C**). While acidification destabilized *β*-sheets, the interaction of salt ions with proteins and solvating water molecules can promote protein folding and stabilization of secondary structure. Upon adding KCl salt to S12-pAzF in water, it retained both *β*-sheet and random coil structures (**Fig. 2C**). When KCl and acetic acid were added simultaneously, we again observed the presence of both *β*-sheet and random coil structures, showing that chloride salt stabilizes *β*-sheets, even in pH conditions that would otherwise be destabilizing.

With the interest of controlling self-assembly, we next asked whether *β*-sheet character could be restored by adding KCl after acidification and *β*-sheet unfolding. Indeed, introducing KCl to acidified S12-pAzF caused chain collapse and re-emergence of *β*-sheets, likely via charge neutralization by chloride ions on positively charged His residues (**Fig. 2C**).^32^ These results demonstrate that *β*-sheets in S12-pAzF can be reversibly unfolded and re-folded, allowing control of chain collapse by manipulating solvent conditions.

Salt anions, which mediate chain collapse and assembly in suckerins have complex interactions with positively-charged His side chains, the protein backbone, and with solvating water molecules.^30,38–40^ Also, choice of salt and ionic strength can impact kinetics of protein NA assembly, and in some cases, the structure and release profile of the resulting particles.^41–43^ The Hofmeister series describes some salts as kosmotropes (derived from Greek for *order*), characterized by a tendency to stabilize, or increase the attractive interactions between hydrophobes in solution.^40^ To characterize differences between four common Hofmeister series kosmotropes (**Fig. 2D**) for salting-out S12-pAzF, we employed a high-throughput spectrophotometric method where optical density at 350 nm (OD 350 nm) was measured, noting that increases in OD are due to the assembly of higher-order structures that scatter light and increase solution turbidity.^44,45^ **Fig. 2E** shows representative photographs of high turbidity (OD>1) and low turbidity (OD=0.1) S12-pAzF samples. Because pH impacts solubility, secondary structure, and charge-screening effects, we tested each salt at a range of pHs.

In citrate and sulfate salts (the most kosmotropic salts), we observed high turbidity (OD>1) over the whole pH range tested, indicating the presence of large aggregates regardless of pH, and little dynamic range for tuning particle size under these conditions (**Fig. 2F**). In the lesser kosmotropic chloride and acetate salts, we observed a pH-dependent response and a higher dynamic range for tuning the assembly state of S12 using pH. Specifically, below pH 4.5, turbidity remained low (OD<0.2), indicating conditions for fabricating nanoscale assembles, or a lack of assembly. Above pH 4.5, but below the pKa of the imidazole side chain of His (pKa ~6), we observed significant increases in OD, indicating assembly via charge screening on His residues (**Fig. 2F**). As the pH approached 7, high turbidity was observed in all samples, indicating that deprotonation of His residues results in unconstrained aggregation with or without salt. Notably, these trends were observed in both sodium and potassium salts, providing evidence that anion-mediated effects dominate the observed behavior (**Fig. S3**).

These results show that S12 salting-out follows the direct Hofmeister series, with the most kosmotropic salts precipitating S12 most efficiently over the conditions tested. This corroborates our previous observations that chain collapse in crosslinked S12 hydrogels was mediated by anions, and that kinetics of hydrogel contraction trended with the kosmotropic arm of the Hofmeister anion series.^30^ We postulate that differences in pH-response across salts is due to (i) differences in anion avidity for charged His residues, (ii) the mediation of interactions with other parts of the protein (such as hydrophobic patches in amorphous domains) by the more kosmotropic salts. Future studies using quantitative methods to discern structure and protein/solvent interactions, such as FTIR or nuclear magnetic resonance spectroscopy, could be used to elucidate the mechanisms behind the observed behavior. Additionally, the use of molecular dynamics has already been employed to study the structure of minimal suckerin peptides, and could be applied to understand in greater depth the molecular mechanisms behind our findings.^26,46^ From a practical perspective, these results show which salts are useful for engineering pH-dependent self-assembly processes with suckerins, and addresses questions about the impact of salt identity on self-assembly.

### Salting-out and functionalizing S12 NAs

With an understanding of the process variables impacting *β*-sheet enrichment and solubility of S12, we sought to characterize S12-pAzF NAs salted out with a chloride salt. In initial optimizations, we used dynamic light scattering (DLS) to characterize influences of protein concentration, acidification, and salt concentration on particle size distributions. **Fig. S4** shows average diameter (Z_ave_), polydispersity index (PDI), and transmission electron microscopy (TEM) characterization of select conditions. Particle size distributions (PSDs) are presented in **Fig. S5.** We observed several trends from these experiments. The factor that contributed most significantly to constraining particle sizes was acidification prior to addition of KCl. These conditions resulted in the formation of discrete NAs with narrow size distributions (**Fig. S5**). When the solvent was not acidified, we observed relatively large average particle diameters, higher PDIs, and the formation of micron-scale, irregularly shaped aggregates of particles upon addition of KCl. We also observed that decreasing protein concentration and lowering the process temperature decreased average particle sizes, suggesting that controlling the kinetics of aggregation constrains particle diameters.

Guided by our optimizations, we identified conditions for fabricating ~100 nm particles, since this is a desirable size regime for biomedical applications. We pre-acidified S12-pAzF, then salted out NAs fabricated at three concentrations of protein using 250 mM KCl. Here, we observed that average particle sizes increased as the protein concentration was increased. We confirmed the rank-order of average particle sizes using two in-solution dynamic light scattering (DLS) techniques: a bulk measurement using a Zeta Sizer, and Nanoparticle Tracking Analysis (NTA), a technique which individually tracks and sizes particles in a flowcell over time. Specifically, for particles fabricated at 0.02, 0.06, and 0.11 mg/mL of S12-pAzF, particle sizes were ~100 nm, ~120 nm, and ~150 nm, respectively, with particle sizes increasing with protein concentration (**Fig. 3A**). PSDs and PDIs measured via Zeta Sizer are shown in **Fig. 3B-3C**, and the associated NTA PSDs are presented in **Fig. S6**. PDI, a metric of the broadness of the measured size distribution that calculated directly from the DLS correlation data, was low (<0.15) for each condition, showing uniform particle sizes. TEM images are shown for the 0.11 mg/mL condition in **Fig. 3D**, confirming the presence of circular ~150 nm particles, consistent with scattering data. Images for the 0.02 and 0.06 mg/mL conditions are presented in **Fig. S6.** The morphologies of NAs indicated that the particles were protein-core, as opposed to hollow-core or micellar. The methods used here to salt-out S12-pAzF NAs are similar to those used previously for S19 NAs, indicating that the experimental techniques used here could be generalizable for characterizing self-assembly of other suckerins, both natural and synthetic.

**Figure 3.**
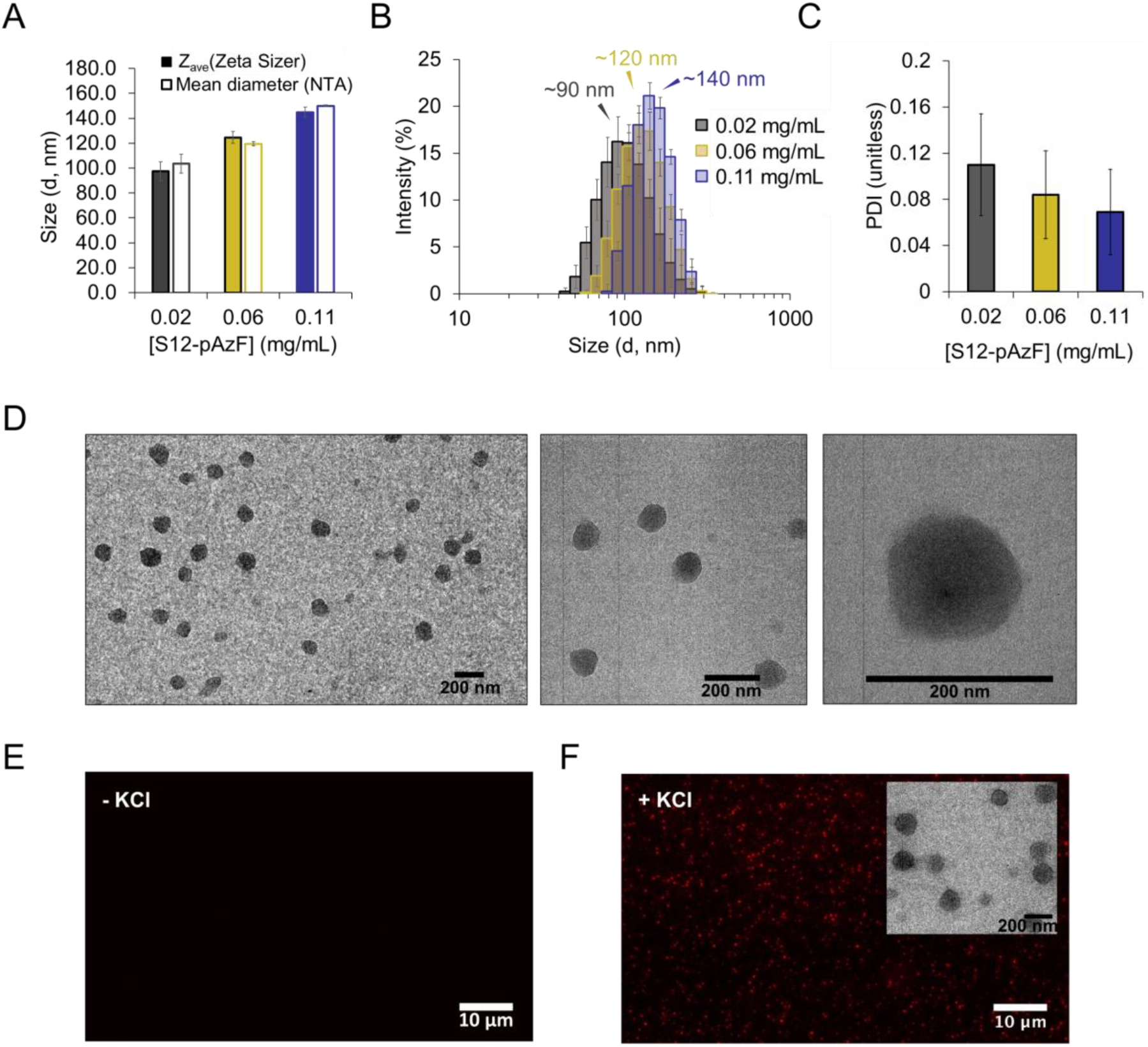
Salting out and functionalizing S12 NAs. **(A)** DLS analysis of S12-pAzF NAs fabricated at three concentrations of S12-pAzF. Solid bars show the average diameter (Z_ave_) measured by Zeta Sizer and outlined bars show the mean diameter measured via Nanoparticle Tracking Analysis (NTA). Error bars represent the standard deviation between measurements of three independently prepared batches of NAs. **(B)** Particle size distributions of S12-pAzF NAs. Annotations indicate the size corresponding with the approximate intensity maximum measured in each sample. Error bars represent the standard deviation between three independently prepared batches of NAs. **(C)** Polydispersity indices of NAs measured by Zeta Sizer. Error bars represent the standard deviation between three independently prepared batches of NAs. **(D)** Representative TEM micrographs of NAs fabricated at 0.11 mg/mL of S12-pAzF, with magnification increasing from left to right. **(E)** Fluorescence microscopy image of S12-TAMRA with no salt added. **(F)** Fluorescence microscopy image of S12-TAMRA with KCl added to salt out NAs. ***Inset*** shows TEM micrograph of the corresponding fluorescent S12-TAMRA NAs. Data are representative of three independent experiments.

Previous efforts for preparing functional self-assembled suckerin NAs involved sequestration of hydrophobic drug cargo through noncovalent interactions.^32^ Noncovalent loading methods, however, suffer from lack of control of the loading density and release of cargo. Covalent conjugation of cargo offers an improved level of control for fabricating drug-loaded therapeutic nanocarriers and fine-tuning release profiles.^47–50^ Toward this eventual goal, we fabricated particles using S12-TAMRA, since the TAMRA fluorophore is a hydrophobic molecule. Using fluorescence microscopy, we observed the formation of dispersed, circular particles only upon the addition of KCl (**Fig. 3E-F**). The morphology and size of TAMRA-conjugated NAs were similar to NAs fabricated without a fluorophore, showing that self-assembly proceeds similarly with a conjugate small molecule (**Fig. 3F, inset**). Since the relatively small size of the conjugated fluorophore (0.9 kDa) allowed self-assembly to proceed unhindered, we hypothesize that small-molecules of similar sizes and hydrophobicities to TAMRA, like small-molecule cancer therapeutics, could likely be loaded into S12 NAs using the same approach. These results pave the way for future studies to fabricate precisely defined and drug loaded NAs. Since multiple installations of pAzF (>30) within a single protein are accessible with the OTS used here, future studies could also interrogate the feasibility of higher small-molecule loading densities.^36,51^

### PEGylation of S12 changes assembly properties

We next sought to leverage site-specific functionalization to rapidly build and test conjugates of S12 with altered hydrophobicity. Altering the compositions of hydrophobic: hydrophilic sequences in polymers is a strategy for engineering self-assembly and crystallinity of block copolymers.^22,52–59^ For proteins, this process can often involve tedious cloning procedures (since highly-repetitive DNA sequences are challenging to synthesize), and protein expression of many variants can be daunting.^60^ Thus, the ability to use SPAAC to controllably functionalize proteins with hydrophilic ligands could expedite designing and building synthetic amphiphiles, and could eventually guide the design of engineered protein sequences with precisely-defined properties.

The amino acid sequence of S12 is mostly hydrophobic (**Fig. 4A**), and the hydrophobic interactions of inter- and intra-molecular *β*-sheets are critical for S12 self-assembly. Thus, we reasoned that covalently conjugating a model hydrophilic polymer, polyethylene glycol (PEG), to S12 would alter protein properties. Using SPAAC, we conjugated S12-pAzF to a 5 kDa PEG (referred to as S12-5K) molecule with >90% efficiency (**Fig. 4B**). We tested for changes in the solubility between S12 and S12-5K in water, and in two buffer/pH combinations where the unconjugated protein is insoluble: MES buffer at pH 6.6, and potassium phosphate buffer at pH 8. PEGylation improved the solubility of S12, by 28% and 11% at pH 6.6 and pH 8, respectively, showing that increasing hydrophilicity of S12 improves solubility at near or above physiological pH (**Fig. 4C**). S12-5K formed circular particles on the order of 30-50 nm in size in water, where unconjugated S12-pAzF did not exhibit measurable particle formation or stable DLS spectra (**Fig. 4D**). Increasing the molecular weight of the PEG molecule to 10K changed the properties further, increasing the solubility of the conjugate to >75% at pH >6, and changing the morphology to a soluble fibrillar network of particles (**Fig. S7**).

**Figure 4.**
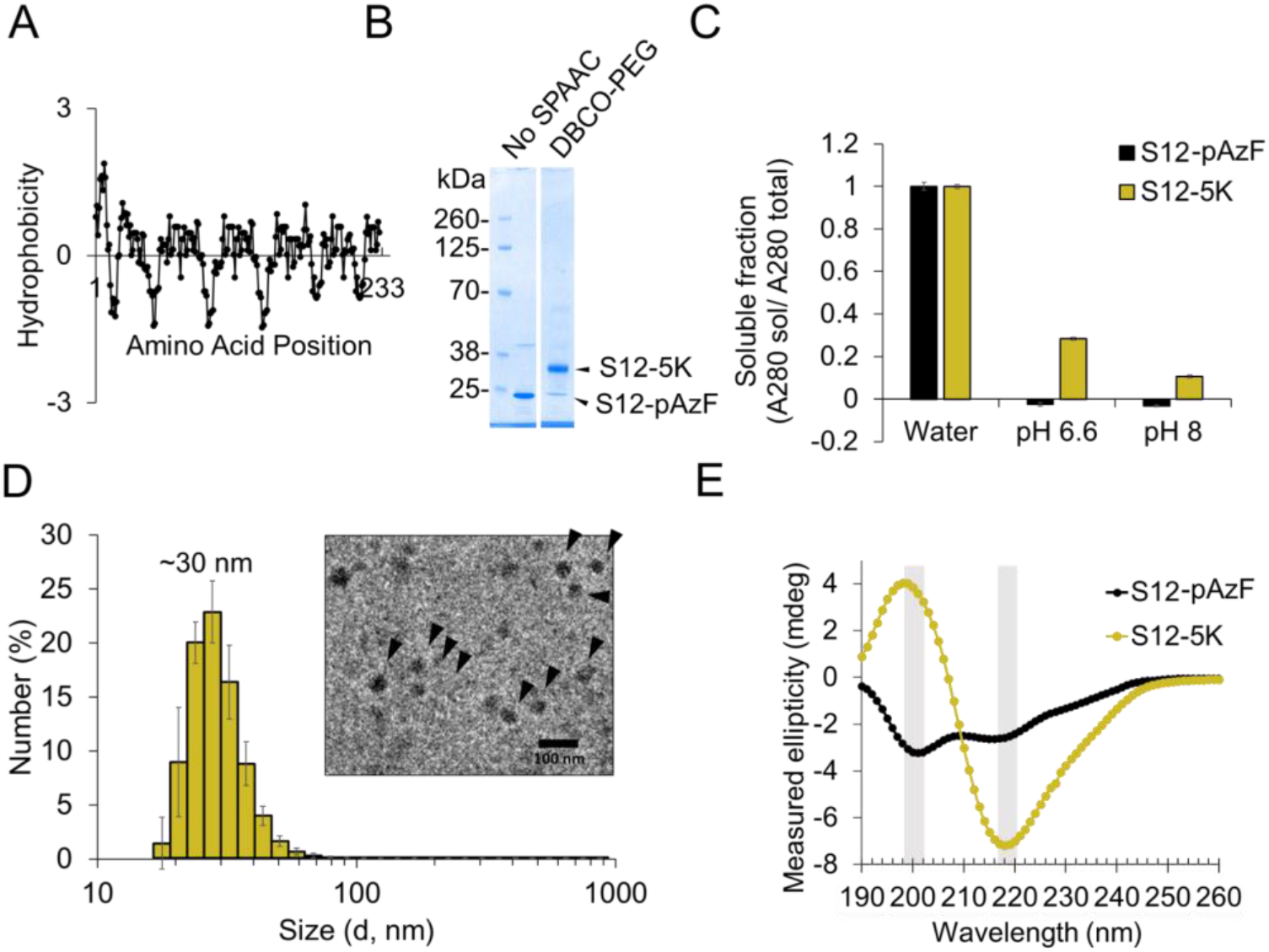
PEGylation of S12 changes assembly properties. **(A)** Kyte & Doolittle hydrophobicity plot of S12. Residues corresponding with positive values are hydrophobic. Plot was generated using the ProtScale (ExPASy) using a window size of 9 and a linear weight variation model. **(B)** SDS-PAGE analysis of S12-5K, showing conjugation of 5 kDa PEG to S12-pAzF via SPAAC. Unconjugated S12-pAzF and the S12-5K conjugation product are indicated. Uncropped gel is shown in **Fig. S7**. **(C)** Soluble (sol) fraction S12-pAzF and S12-5K in water, MES pH 6.6, and Tris pH 8. Error bars represent the standard deviation of three independently prepared samples. **(D)** DLS analysis of S12-5K in water. ***Inset*** shows an electron micrograph of S12-5K particles with black arrows indicating particles for clarity. Error bars represent the standard deviation of three independently prepared samples. **(E)** CD analysis of S12-pAzF and S12-5K in water. Gray rectangles highlight the wavelength ranges corresponding with random coil (~200 nm) and beta sheet (~217-220 nm) signatures. Traces represent the average of triplicate background subtracted scans.

To better understand the impacts of PEGylation on S12-pAzF properties, we studied the secondary structure of S12-5K. Despite its increased hydrophilicity, S12-5K displayed a significant enrichment in *β*-sheet character compared with S12-pAzF, potentially increasing its propensity to self-assemble. The CD spectra of both S12-pAzF and S12-5K displayed a broad minimum near 210-220 nm (indicating *β*-sheet character), with the minimum being more pronounced for S12-5K. However, where S12-pAzF displayed a minimum at 200-210 nm, indicative of random coil, the S12-5K spectrum displayed a switch to positive ellipticity with a peak between 200-210 nm. These spectral features indicate a significant decrease in random coil signature and an enrichment of *β*-sheet content in S12-5K. Taken together, our results indicate that PEGylation, while increasing the solubility of S12 by increasing hydrophilicity, also induced changes in *β*-sheet content, orientation, and presentation in S12-5K. Since *β*-sheet formation is a driving force for molecular self-assembly, modulation of *β*-sheet content, and therefore PEGylation, changes the propensity for inter- and intra-protein hydrophobic interactions. While further studies are needed to understand the impact of PEG length on secondary structure and emergent properties, our results demonstrate that the alteration of S12-pAzF using PEGylation via SPAAC can result in conjugates that have altered solubility and assembly properties.

## Conclusions

Here, we characterized the salt and pH responsiveness of suckerin-12, a promising protein for fabricating functional NAs. We optimized self-assembly and identified conditions for fabricating ~100 nm suckerin-based NAs using a simple salting-out procedure. We anticipate that our workflow will be a useful framework for future efforts in fabricating and processing natural and synthetic suckerin materials. Finally, we showed that the ncAA pAzF is useful for obtaining high yields of recombinant S12-pAzF, which can be efficiently functionalized using SPAAC. The ability to site-specifically modify S12 allowed us to fabricate ~100 nm fluorophore-loaded NAs, and to alter the assembly properties of S12. In future studies, the S12-pAzF scaffold described here could be used to make functional materials such as NAs with precise drug-loading or self-assembling amphiphiles where functional hydrophilic molecules other than PEG (such as DNA or carbohydrates) are used to impact assembly and enhance functionality. Ultimately, we anticipate that ncAAs will expedite the development of functional nanomaterials by allowing researchers to quickly synthesize new protein conjugates with desired properties.

## Associated content/ supporting information

Figure S1. Optimized inclusion body preparation of S12; Figure S2. SPAAC reactions proceed in water and in 1% acetic acid; Figure S3. pH-dependence of S12 solubility in various salts is anion-dependent; Figure S4. Optimization of NA fabrication conditions; Figure S5. Optimization of NA fabrication conditions, continued; Figure S6. Further characterization of NAs fabricated with optimized conditions; Figure S7. Characterization of S12-PEG conjugates; Supporting Information methods: CryoTEM.

## Author contributions

The manuscript was written through contributions of all authors. All authors have given approval to the final version of the manuscript.

## Acknowledgements

We thank the Center of Excellence for Bioprogrammable Nanomaterials (C-ABN) through the Air Force Research Labs for providing funding (FA8650-15-2-5518) and a collaborative infrastructure for our research. This work made use of the EPIC, Keck-II, IMSERC, and facilities of Northwestern University’s NU*ANCE* Center, which have received support from the Soft and Hybrid Nanotechnology Experimental (SHyNE) Resource (NSF NNCI-1542205); the MRSEC program (NSF DMR-1121262) at the Materials Research Center; the International Institute for Nanotechnology (IIN); the Keck Foundation; and the State of Illinois, through the IIN. We thank Miles Markmann and Eric Roth for helpful conversations about TEM. We thank Marquise Crosby for helpful conversations about protein purification. JMH was supported by the National Defense Science and Engineering Graduate Fellowship, the C-ABN, and the Ryan Fellowship. MCJ acknowledges the David and Lucile Packard Foundation and the Camille Dreyfus Teacher-Scholar Program for support. PBD is adjunct faculty at Wright State University, Dayton, Ohio. RRN acknowledges funding support from AFOSR.

## Abbreviations

CD: circular dichroism
DBCO: dibenzocyclooctyne
DLS: dynamic light scattering
FTIR: Fourier-transform infrared
MRE: mean residue ellipticity
NA: nanoassemblies
ncAA: non-canonical amino acids
NTA: nanoparticle tracking analysis
OD: optical density
OTS: orthogonal translation system
pAzF: para*-L*-azido-phenylalanine
PDI: polydispersity index
PEG: polyethylene glycol
PSD: particle size distribution
S12: suckerin-12
S19: suckerin-19
SPAAC: strain-promoted azide-alkyne cycloaddition
SRT: sucker ring teeth
TEM: transmission electron microscopy

