## Supporting Information for "Characterizing and controlling nanoscale self-assembly of suckerin-12"

Contents:

Figure S1. Optimized inclusion body preparation of S12

Figure S2. SPAAC reactions proceed in water and in 1% acetic acid

Figure S3. pH-dependence of S12 solubility in various salts is anion-dependent

Figure S4. Optimization of NA fabrication conditions

Figure S5. Optimization of NA fabrication conditions, continued

Figure S6. Further characterization of NAs fabricated with optimized conditions

Figure S7. Characterization of S12-PEG conjugates

Supporting Information methods: CryoTEM

A

| Well Number | Well Contents | Dilution Factor |
|---|---|---|
| 1 | Standard | - |
| 2 | Whole Cell Resuspension (Pre-lysis) | 10 |
| 3 | Post-lysis Supernatant | - |
| 4 | Triton X-100 Wash 1 Resuspension | 10 |
| 5 | Triton X-100 Wash 1 Supernatant | - |
| 6 | Triton X-100 Wash 2 Resuspension | 10 |
| 7 | Triton X-100 Wash 2 Supernatant | - |
| 8 | Triton X-100 Wash 3 Resuspension | 10 |
| 9 | Triton X-100 Wash 3 Supernatant | - |
| 10 | No Triton Wash Resuspension | 10 |
| 11 | No Triton Wash Supernatant | - |
| 12 | Water Wash Resuspension | 10 |
| 13 | Solubilized Suckerin-12 (Post-high speed spin) | 10 |
| 14 | Pre-dialysis Product | 10 |

B

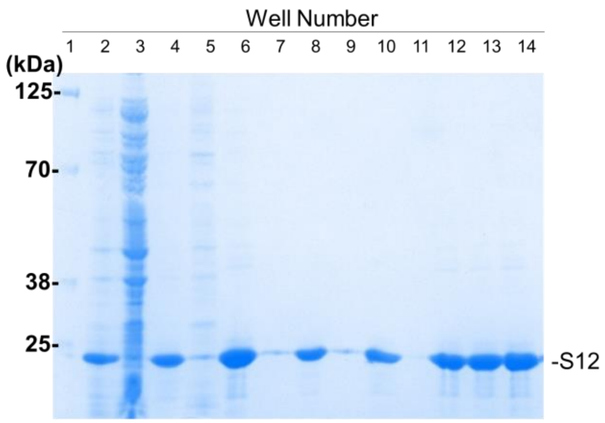


C

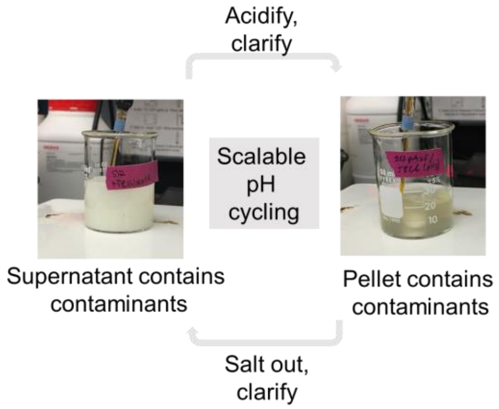


**Figure S1.** Optimized inclusion body preparation of S12. **(A)** Well key of SDS-PAGE gel shown in **(B)**. **(B)** SDS-PAGE of various purification fractions, showing soluble fractions of each step. **(C)** Photographs of precipitated and solubilized S12 from inclusion bodies, showing the concept of scalable 'pH cycling' for purification of S12.

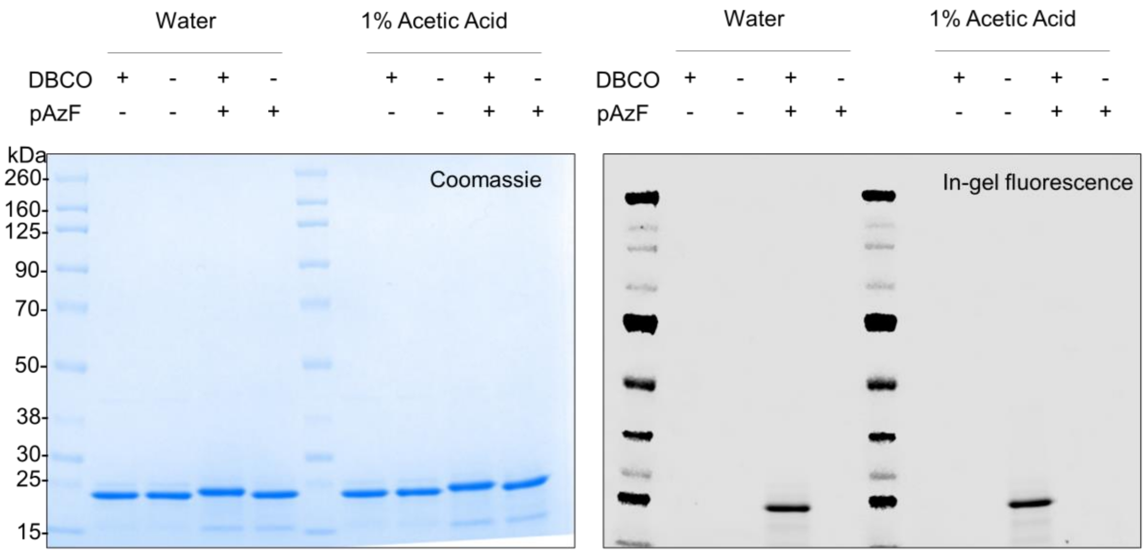


**Fig S2.** SPAAC reactions proceed in water and in 1% acetic acid. Coomassie ***(left)*** and in-gel fluorescence ***(right)*** images of SPAAC reaction products. Reactions were run in water or acetic acid as indicated. -/+ pAzF refers to the protein added to the reaction (S12wt or S12-pAzF, respectively), and -/+ DBCO refers to the absence or presence of the DBCO-TAMRA probe.

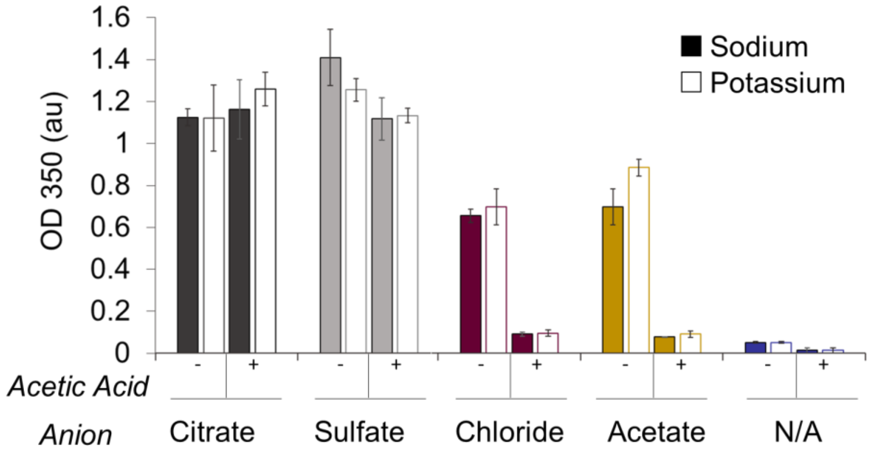


**Figure S3.** pH-dependence of S12 solubility in various salts is anion-dependent. Turbidity measurements of S12-pAzF in sodium and potassium salts of various anions (with or without 5% acetic acid added, indicated with +/-).

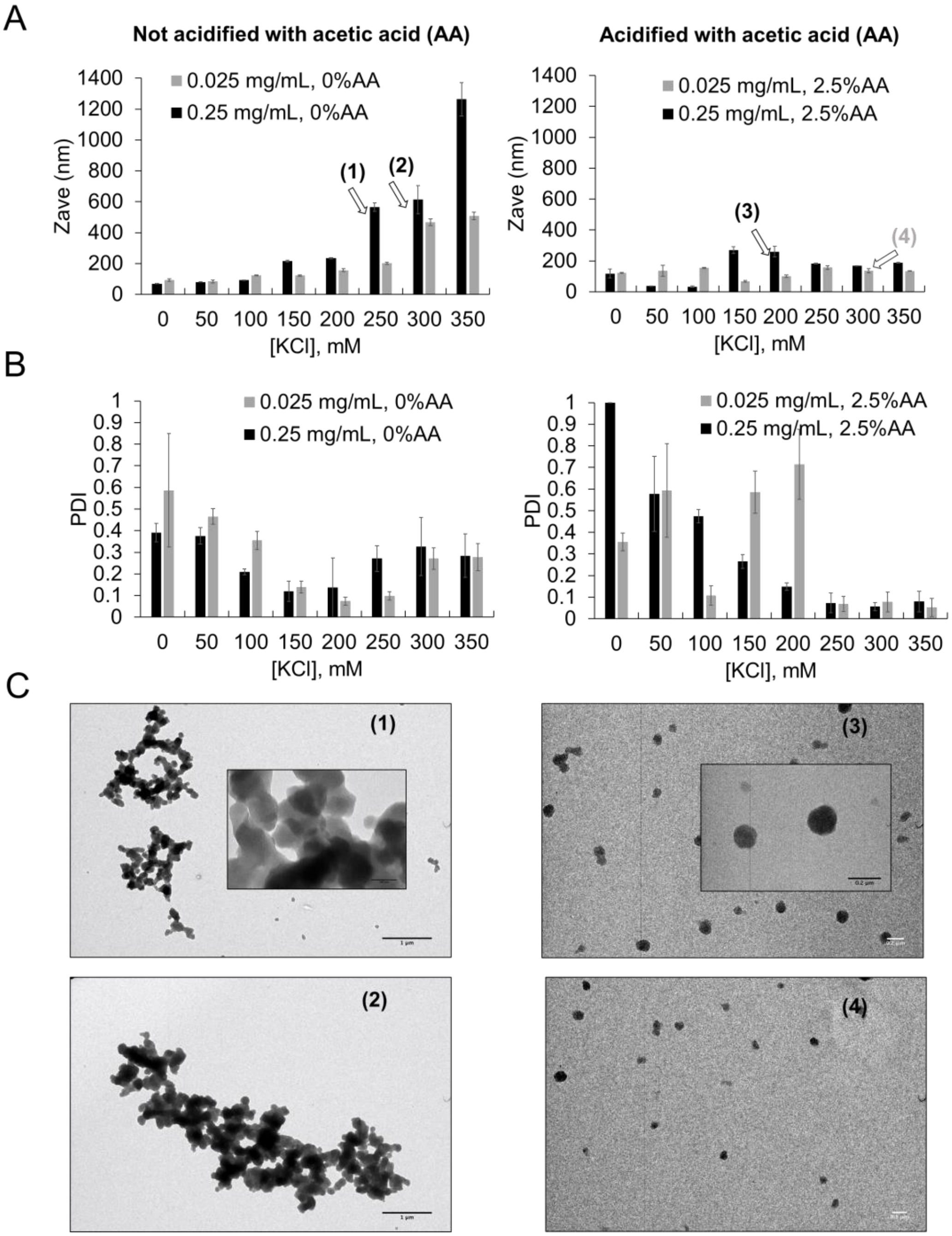


**Figure S4.** Optimization of NA fabrication conditions. **(A)** Average diameters (Zave) of S12-pAzF particles fabricated at the indicated conditions. DLS was conducted using Zeta Sizer. Error bars represent the standard deviation of three measurements. **(B)** Associated polydispercity indexes (PDI) of the samples in **(A)**. Error bars represent the standard deviation of three measurements. **(C)** Selected TEM images of 4 conditions, indicated via numbering and arrows indicating NA fabrication conditions in **(A)**.

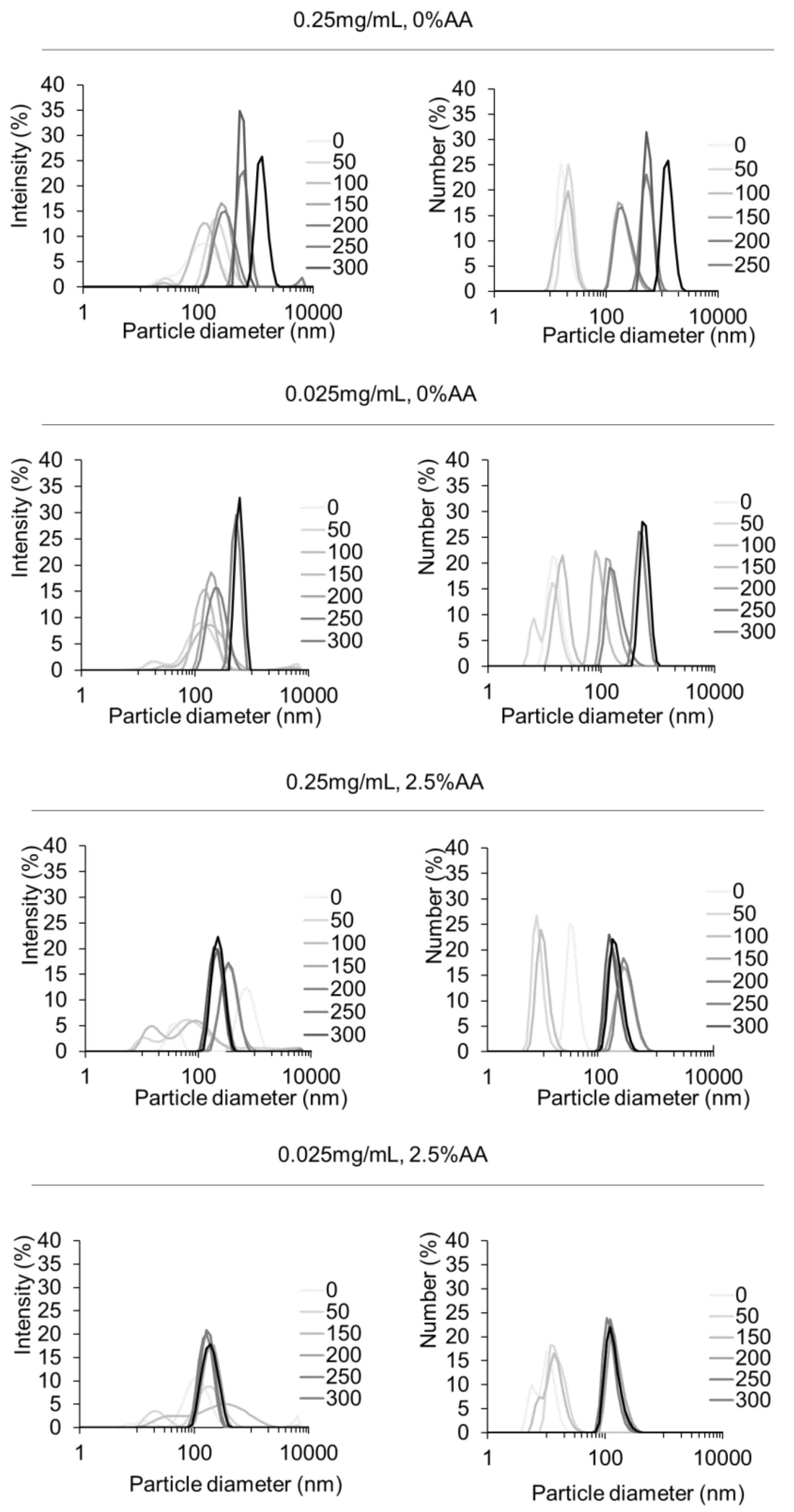


**Figure S5.** Optimization of NA fabrication conditions, continued. Intensity and number particle size distributions of S12-pAzF particles fabricated at the indicated conditions, measured via Zeta Sizer.

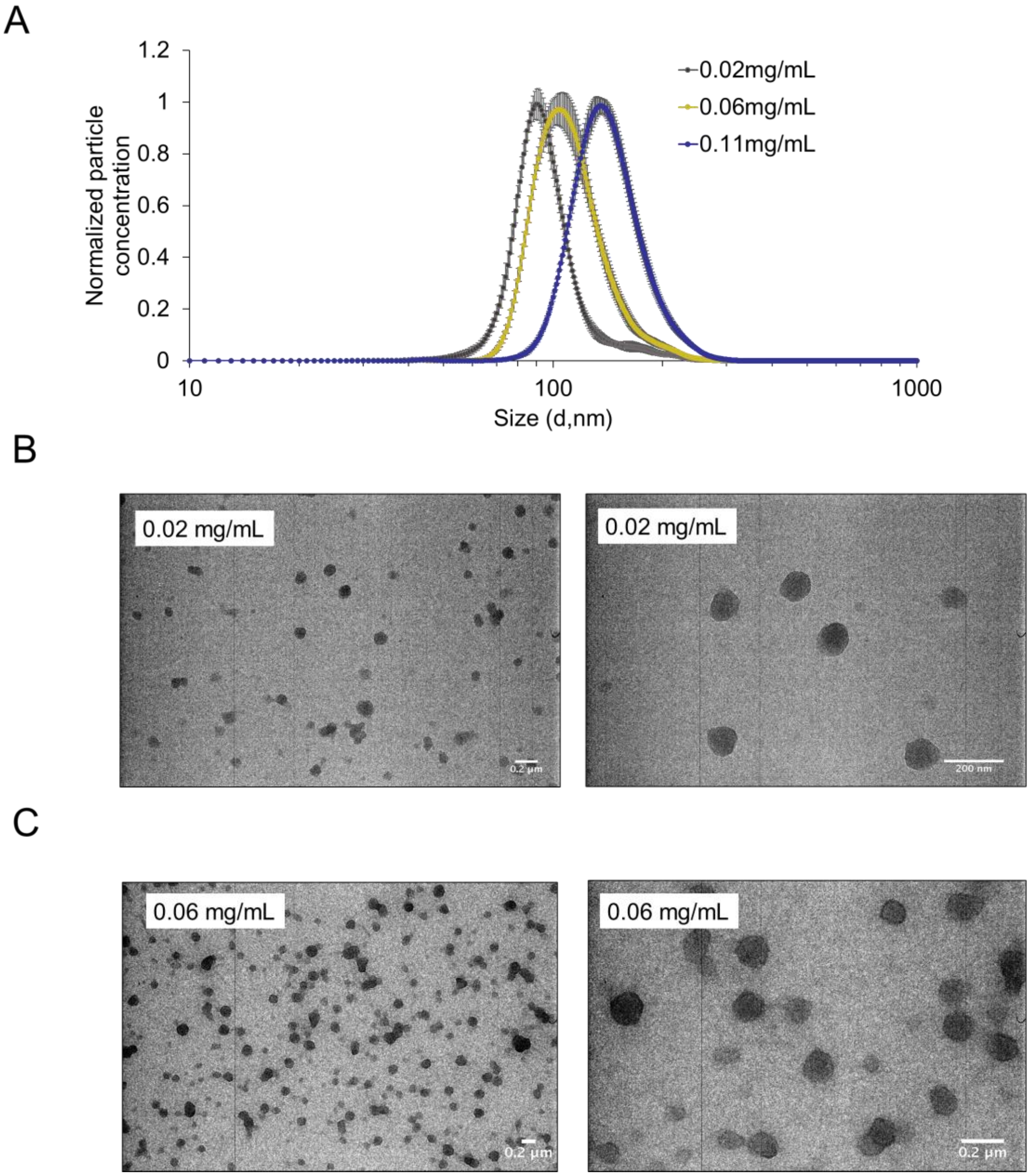


**Figure S6.** Further characterization of NAs fabricated with optimized conditions. **(A)** Particle size distributions obtained via NTA. Error bars represent the standard deviation of measurements of three independently prepared batches of NAs. Associated TEM images for NAs made with 0.02 mg/mL **(B)** and 0.06 mg/mL **(C)** of S12-pAzF.

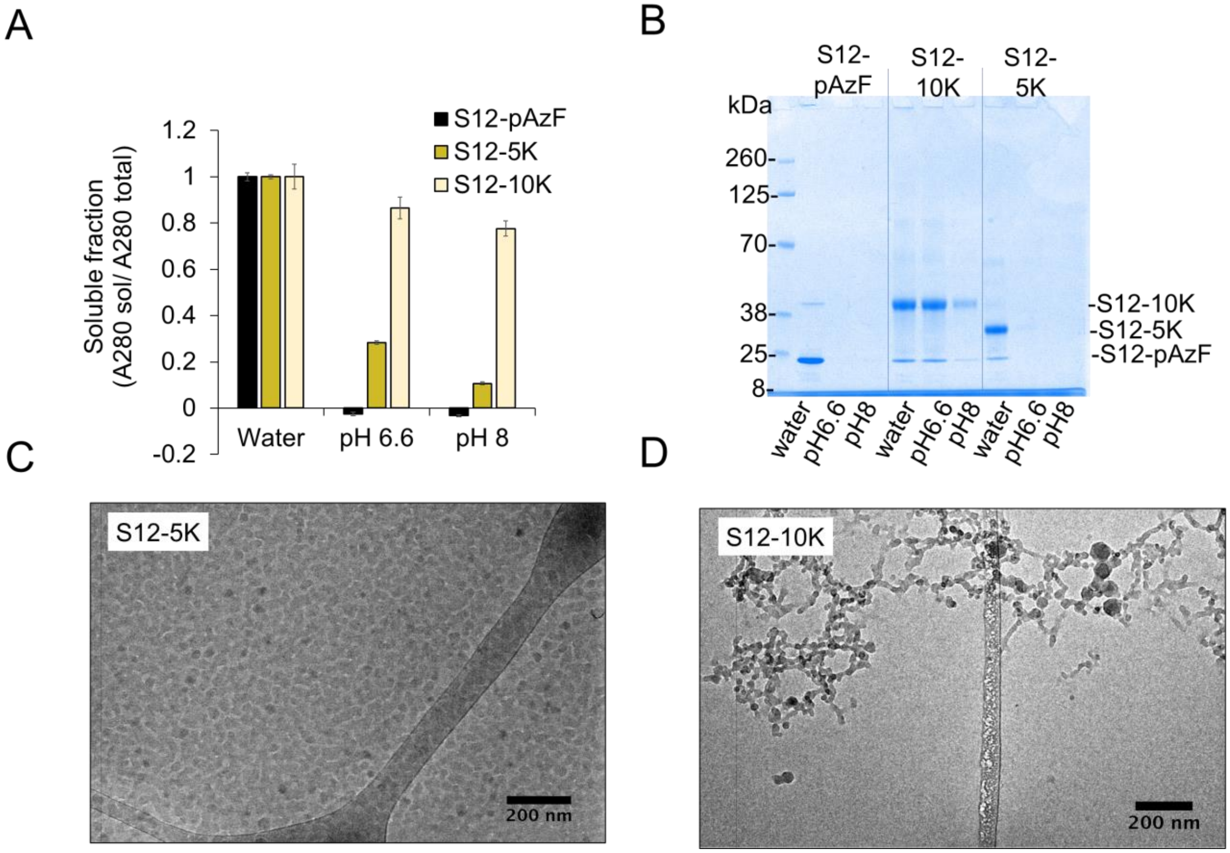


**Figure S7.** Characterization of S12-PEG conjugates **(A)** Soluble (sol) fraction of S12-pAzF, S12-5K, and S12-10K in water, MES pH 6.6, and Tris pH 8. **(B)** SDS-PAGE analysis of the soluble fractions shown in panel A. Annotations indicate which protein construct was added to each well and the associated solvent conditions. CryoTEM images of S12-5K **(C)** and S12-10K **(D)** in water.

**Supporting Information Methods:**

Cryo-TEM was used to analyze the morphologies of S12-5K and S12-10K PEG conjugates. For cryo-TEM measurement, 200 mesh Cu grids with a lacey carbon membrane (EMS Cat. # LC200-CU) were glow discharged as described above. 4 µL of sample at 0.5 mg/mL S12-PEG were pipetted onto the grid and blotted for 5 seconds with a blot offset of +0.5 mm, followed by immediate plunging into liquid ethane within a FEI Vitrobot Mark III plunge freezing instrument (Thermo Fisher Scientific). The plunge-frozen grids were kept vitreous at –172 $^{o}$C in a Gatan Cryo Transfer Holder model 626.6 (Gatan Inc., Pleasanton, CA, USA) while viewing in a JEOL

JEM1230 LaB6 emission TEM (JEOL USA, Inc., Peabody, MA,) at 120 keV. Image data was collected by a Gatan Orius SC1000 CCD camera Model 831 (Gatan Inc.). Image analysis was done using Image J.
